# Increased protein kinase Mζ expression by Minocycline and N-acetylcysteine restores late-phase long-term potentiation and spatial learning after closed head injury in mice

**DOI:** 10.1101/2024.09.20.613738

**Authors:** Elena Nikulina, Panayiotis Tsokas, Kristen Whitney, Andrew Tcherepanov, Changchi Hsieh, Todd C. Sacktor, Peter J. Bergold

**Author notes:** **Corresponding author:** Peter J. Bergold, Ph.D., Department of Physiology and Pharmacology, State University of New York Downstate Health Sciences University, Brooklyn, New York, 450 Clarkson Avenue, Brooklyn, New York11203. Department of Pathology, Icahn School of Medicine at Mount Sinai, New York, New York (Kristen Whitney). **Author contributions** Elena Nikulina, Conceptualization, Investigation, Writing - Review & Editing; Panayiotis Tsokas, Methodology, Investigation; Kristen Whitney, Investigation; Andrew Tcherepanov, Investigation; Changchi Hsieh, Formal analysis; Todd C. Sacktor, Conceptualization, Writing - Review & Editing, Funding acquisition; Peter J. Bergold, Conceptualization, Writing - Original Draft, Project administration, Funding acquisition.

## Abstract

Cognitive deficits frequently arise after traumatic brain injury. The murine closed head injury (CHI) models these deficits since injured mice cannot acquire Barnes maze. Dosing of minocycline plus N-acetylcysteine beginning 12 hours post-CHI (MN12) restores Barnes maze acquisition by an unknown mechanism. Increased hippocampal synaptic efficacy is needed to acquire Barnes maze, synaptic long-term potentiation (LTP) models this increased synaptic efficacy *in vitro*. LTP has an early phase (E-LTP) lasting up to one hour that is mediated by second messengers that is followed by a late phase (L-LTP) that needs new synthesis of protein kinase M zeta (PKMζ). PKMζ has constitutive kinase activity because it lacks the autoinhibitory regulatory domain found in other PKCs. Due to its constitutive activity, the amount of PKMζ kinase activity is determined by PKMζ protein levels. We report that CHI bilaterally decreases PKMζ levels in the CA3 and CA1 hippocampus. MN12 increases CA1 PKMζ expression. CHI inhibits E-LTP in slices from the ipsilesional hippocampus and inhibits L-LTP in slices from both hippocamppi. MN12 treatment reestablishes both E-LTP and L-LTP in slices from the injured MN12-treated hippocampus. The restoration of L-LTP from injured MN12-treated hippocampus is mediated by PKMζ because L-LTP is blocked by the specific PKMζ inhibitor, ζ-stat. Hippocampal ζ-stat infusions also prevents Barnes maze acquisition in injured, MN12-treated mice. These data suggest that post-injury minocycline plus N-acetylcysteine targets PKMζ to improve synaptic plasticity and cognition in mice with closed-head injury.

## Introduction

A closed head injury (CHI) impairs Barnes Maze acquisition that is restored by MN12 treatment (1). Learning Barnes maze requires potentiation of hippocampal synapses that is modeled *in vitro* by long-term potentiation (LTP) (2-9). LTP has a second messenger-mediated early phase (E-LTP) lasting approximately 60 minutes that is followed by a persistent, protein synthesis-dependent late phase (L-LTP) (10). Many rodent TBI models impair E-LTP (11-17) and prevent acquisition of hippocampus-dependent tasks (12-14). Drugs can restore E-LTP in injured rodents (12, 13, 18). No studies have examined if experimental TBI impairs L-LTP.

The protein kinase C isoform PKMζ is a key mediator of L-LTP maintenance. During L-LTP, PKMζ binds the postsynaptic scaffolding protein KIBRA that stabilizes PKMζ (10, 19). The small molecule inhibitor ζ-stat prevents PKMζ-KIBRA binding leading to increased PKMζ turnover and reversal of pre-established L-LTP (19). ζ-stat specifically inhibits KIBRA-PKMζ binding (IC_50_ = ∼1 µM) and 10-fold higher ζ-stat did not prevent KIBRA binding to the protein kinase C isoform, PKCι*/*λ that has high homology to PKMζ (19).

This study examines how the drug combination of minocycline (MINO) and N-acetylcysteine (NAC) restores L-LTP and Barnes maze acquisition. At concentrations used in this study, MINO is anti-inflammatory, anti-apoptotic; chelates iron and inhibits metalloprotease 9 (20). NAC is an anti-oxidant that elevates extracellular glutamate levels through the X_c_ cysteine-glutamate antiporter (20). This study examines if MINO plus NAC improves cognition after CHI by targeting PKMζ expression.

## Methods

### Closed Head Injury and drug treatments

Closed head injury is done as described by Sangobowale, *et al*. (1). Briefly, isoflurane anesthesia is induced (3.5% in O_2_, 1.0 L/min) in male C57/BL6 mice (15-17 weeks, 26-28gr, Jackson Laboratories) and maintained (3% in O_2_ (1.0 L/min) until completion of the head impact. An electromagnetically controlled piston (Leica Microsystems, Buffalo Grove, IL) impacts the mouse head above the left parietal lobe (1). Sham-injured mice receive the same treatment without the head impact. Time to regain righting reflex is measured immediately after head impact. Mice with a righting reflex under 330 seconds are excluded from the study. Mice are randomly assigned into a CHI group treated with saline or MN12 (MINO, 22.5 mg/kg; NAC 75 mg/kg) or a sham-CHI group treated with saline. Saline or MN12 is administered intraperitoneally at 12 hours, 1 day, and 2 days after Sham-CHI or CHI. Drugs are from Millipore-Sigma (St. Louis, MO).

### Electrophysiology

Hippocampal slices are prepared and recorded as described by Tsokas, *et al*. (21). Slices are eliminated if field excitatory post-synaptic potential (fEPSP) spike threshold is <2 mV. WinLTP data acquisition program analyzes the maximum slope of the rise of the fEPSP. Baseline fEPSP slope is assessed for 30 minutes by stimulating the Schaffer collateral/commissural once every 30 seconds. LTP is induced with a high-frequency stimulation (HFS) of two 100 Hz 1-second tetanic trains at 25% of spike threshold, spaced 20 sec apart. E-LTP is assessed at 45 minutes post-HFS, L-LTP is assessed at 165 minutes post-HFS. Slices are treated 90 minutes after HFS with phosphate-buffered saline (PBS) or ζ-stat (10 µM, 1-naphthol-3,6,8-trisulphonic acid, NSC 37044, Drug Synthesis and Chemistry Branch, Developmental Therapeutics Program, Division of Cancer Treatment and Diagnosis, National Cancer Institute) in PBS.

### Barnes maze

Barnes maze is done as described by Sangobowale *et al*. (1). For four days, mice receive four 3-min training trials with a 15-min intertrial interval. An observer blind to the experimental treatment analyzes videos of each training session using AnyMaze software (Stoelting). Latency, and the number of erroneous holes searched is measured across the training trials. Speed is measured on trial day 1. Barnes maze is acquired if latency or errors on training day 4 are significantly lower than on training day 1.

### Histology

Brains from Isoflurane (3–5%, 0.8 L/min in O_2_) anesthetized mice are fixed transcardially with paraformaldehyde (4%, w/v). Parasagittal brain sections (5 µm) from the ipsilesional and contralesional hemispheres (HistoWiz, Inc., Brooklyn, NY) are stained with anti-PKMζ antibodies (21). NIH Image J software assesses PKMζ immunoreactivity in CA3 (from bregma [in mm]) AP, 1.1-1.6; L, +/-0.9-1.4; DV, 1.3-1.8) and CA1 (from bregma [in mm] AP, 1.1-1.6; L, +/-0.9-1.4; DV, 1.0-1.3) (22, 23).

### ζ-stat hippocampal infusions

Mice are treated with carprofen (5 mg/ml), ketamine (1 ml/100g) (10 mg/ml) and xylazine (2 mg/ml), and stereotaxic cannulas implanted (Plastics One, Roanoke, VA) into both dorsal hippocamppi (from bregma [in mm] AP, -2.0 mm; L, +/-0.5 mm; DV, -1.6 mm) as described by Garcia-Osta, *et al*. (24). After a one-week, mice are infused with ζ-stat (6 nmol in PBS, pH 7.4 in 0.3 µl) or PBS using microinjectors (28-g) joined to Hamilton syringes. Mice are infused 30 minutes prior to each day of Barnes maze training. At the end of Barnes maze training, injection cannula location is determined.

### Statistics

Two-way ANOVA analyzes hippocampal PKMζ expression with pairwise comparisons analyzed by Sidak’s post-hoc test. Hippocampal EPSPs are analyzed by repeated value ANOVA at 5 min pre-HFS, or at 45 or 165 minutes post-HFS. ζ-stat effects on hippocampal EPSPs are analyzed by repeated value two-way ANOVA 330 minutes after ζ stat administration with pairwise comparisons by Student-Neuman Keul’s *post-hoc* test. Repeated value two-way ANOVA analyzes Barnes maze latency and error with pairwise comparisons made using uncorrected Fisher’s least significant difference *post-hoc* test. One-way ANOVA analyzes speed on Barnes maze. All values are presented as mean ± standard error of the mean.

## Results

### MN12 prevents CHI-dependent hippocampal loss of PKMζ

PKMζ activity is needed to acquire hippocampus-dependent spatial tasks such as Barnes maze (25). Barnes maze acquisition is impaired in CHI-injured mice and is restored by MN12-treatment (1). This suggests that injury may lower PKMζ expression that is increased by MN12. We tested this hypothesis by examining hippocampal PKMζ expression in injured mice treated with MN12 or saline at 14 days post-injury (DPI) (Figure 1A, B). PKMζ expression in CA3 and CA1 have a significant effect of treatment (F_1,30_= 9.3, p=0.005), and region (F_4,30_= 9.8, p<0.0001), but not interaction of treatment and region (treatment*region) (F_4,30_=2.4, p=0.07). CHI lowers CA3 PKMζ immunoreactivity in the ipsilesional hippocampus (*p<0.05) that is unaffected by MN12 treatment. In contrast, CHI bilaterally lowers CA1 PKMζ immunoreactivity (ipsilesional, p<0.005, contralesional, p<0.005) that is increased by MN12 (ipsilesional, p=0.01, contralesional, p<0.05). These data suggest that MN12 restores hippocampal PKMζ expression in CA1, but not CA3 (Figure 1A, B).

**Figure 1.**
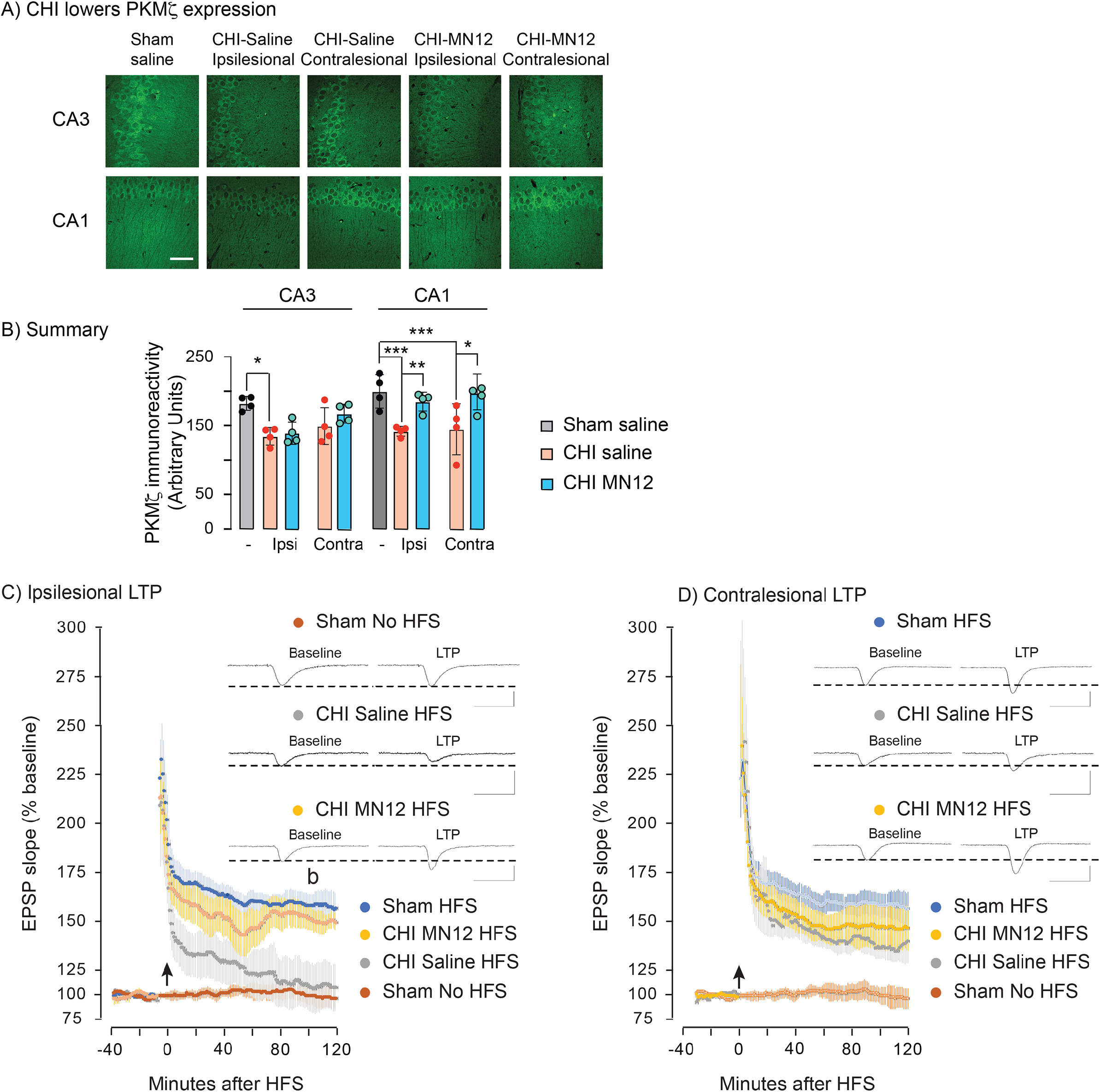
MN12 restores hippocampal PKMζ expression and L-LTP in injured mice. **Panel A**, Representative images of hippocampal CA3 and CA1 PKMζ immunoreactivity. Scale bar, 50µm. **Panel B**, Summary of changes in PKMζ expression. PKMζ expression in CA3 and CA1 has a significant effect of treatment (F_1,30_=9.3, p=0.005), hemisphere (ipsilesional vs. contralesional, F_4,30_= 9.8, p<0.0001), without an interaction of treatment and hemisphere (treatment*hemisphere) (F_4,30_=2.4, p=0.07). CHI lowers CA3 ipsilesional PKMζ expression (*p<0.05). CHI lowers CA1 PKMζ expression bilaterally (***p<0.005) MN12 significantly increases CA1 PKMζ expression (ipsilesional, **p=0.01, contralesional, *p<0.05). Sham (n=5), CHI saline-treated (n=5), CHI MN12-treated (n=6) are analyzed. Changes in fEPSP slope following HFS (arrow) in the ipsilesional (Panel C) or contralesional hippocampus (Panel D). There is a significant effect of time (45 min vs 165 min post-HFS, *F*_1,6_ =7.50, p= 0.03), but not hemisphere (ipsilesional vs. contralesional, F_1,6_=0.19, p=0.7), or time*hemisphere (F_1,6_=1.93, p=0.21). These data suggest disruption of both E-LTP and L-LTP. For MN12 treatment, there is a significant effect of time (5 min pre-HFS vs 165 min post-HFS, F_1,6_=26.0, p=0.002), and no effect of hemisphere (ipsilesional vs. contralesional, F_1,6_=0.20, p=0.7), or time*hemisphere (F_1,6_ = 0.20, p= 0.7). MN12 treatment restored both E-LTP (ipsilesional, p=0.001; contralesional, p=0.001) and L-LTP (ipsilesional, p=0.003; contralesional, p=0.03). An arrow indicates the time of the high frequency stimulation. Representative EPSP traces are shown at baseline (a) and during L-LTP (b). fEPSP slopes in the ipsilesional and contralesional hippocampus are analyzed as a single group, but separate panels present fEPSPs from each hippocampus to better illustrate the individual groups. The same sham saline-treated group with or without HFS is present in panels C and D to assist comparison with injured groups. (n=4 for all groups). Scale Bar; y-axis 1mV, x-axis 20 msec.

### MN12 restores LTP

Treatments that block LTP prevent Barnes maze acquisition (2, 3). Injured mice do not acquire Barnes maze, suggesting that CHI impairs LTP (2-9). LTP is assessed in *ex vivo* slices from sham mice treated with saline or injured mice treated with saline or MN12. LTP is evaluated in both the ipsilesional and contralesional hippocampus. fEPSPs are assayed at baseline (5 minutes pre-HFS), or at 45 minutes (E-LTP) or 165 minutes (L-LTP) post-HFS (Figure 1C). Slices from the ipsilesional saline-treated injured hippocampus lack E-LTP (p=0.14) or L-LTP (p=1.0). Slices from contralesional saline-treated hippocampus express E-LTP (p=0.004), but not L-LTP (p=0.07). MN12 treatment restores both E-LTP (ipsilesional, p=0.001; contralesional, p=0.001) and L-LTP (ipsilesional, p=0.003 contralesional, p=0.03).

MN12 may restore L-LTP by increasing PKMζ expression (Figure 1A,B). The specific PKMζ inhibitor, ζ-stat, inhibits L-LTP after E-LTP is established (19). Ipsilesional slices from injured MN12-treated mice are treated with ζ-stat to test whether MN12 restores L-LTP by targeting PKMζ. At 90 minutes post-HFS, ζ-stat or PBS is bath-applied and fEPSPs recorded for an additional 240 minutes (Figure 2A). fEPSP slope at 330 minutes post-HFS have a significant effect of treatment (F_1,6_=11.7, p=0.01), time (F_2,12_=88.9, p<0.0001), and treatment*time (F_2,12_=6.2, p= 0.01). At 330 minutes post-HFS, ζ-stat, but not PBS, inhibited L-LTP (p= 0.01). These data suggest that MN12 restores L-LTP by increasing PKMζ expression in the injured hippocampus (Figure 2A).

**Figure 2.**
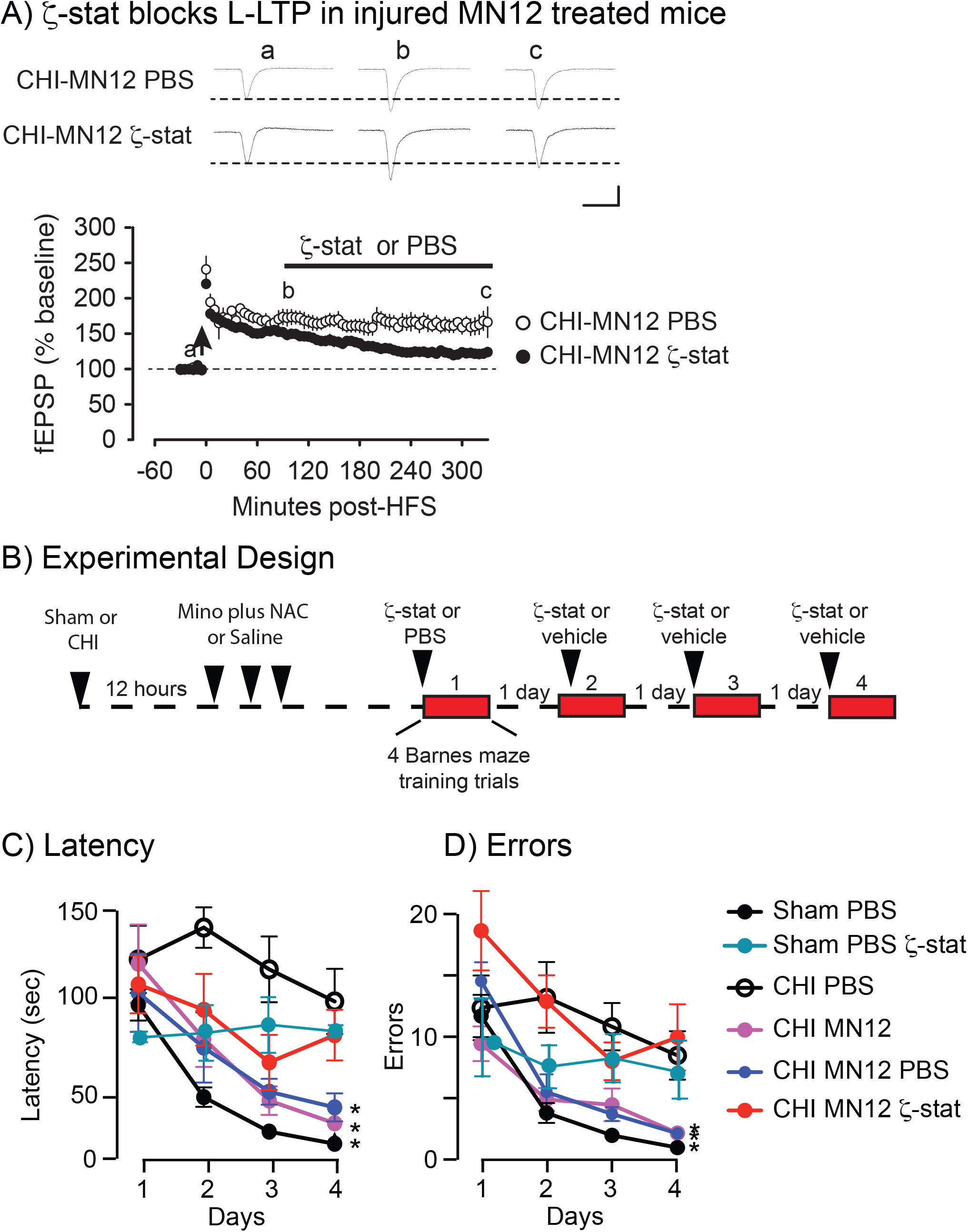
PKMζ is needed for the restoration of L-LTP and spatial learning by MN12. **Panel A, ζ-stat inhibits L-LTP in MN12 treated mice** (n=4 for both groups). Significant effects are seen with treatment (F_1,6_=11.7, p=0.01), time (F_2,12_=88.9, p< 0.0001), and treatment X time (F_2,12_=6.2, p=0.01). At 3 hours after the onset of ζ-stat treatment, fEPSP slope was significantly lower in ζ-stat-treated slices with compared to PBS-treated slices (p**=0.01) (n=4 for all groups). Scale Bar; y-axis 1mV, x-axis 20 msec. **Panel B**, Experimental Design used to test whether PKMζ is needed for restoration of Barnes maze acquisition by MN12. Twelve hours after sham or CHI-injury, mice received a first dose of MN12 or saline treatment followed by daily doses at 2 and 3 DPI. Between 14 and 18 DPI, mice received daily Barnes maze testing. Thirty minutes before each training block, the dorsal hippocamppi are infused bilaterally with either ζ-stat or PBS. **Panel C**, Latency to find the escape hole has a significant effect of days (F_2.0,57.9_=29.6, p<0.0001), treatment (F_5,28_=8.4, p<0.0001) but not days X treatment (F_15,84_=1.4, p=0.17). On day 4, sham PBS treated, CHI-MN12 treated and CHI-MN12 PBS infused mice significantly lowered time to find the escape hole from day 1 (*p<0.05). **Panel D**, Number of errors prior to finding the escape hole has a significant effect of days (F_2.0, 50.5_=22.0, p<0.0001), treatment (F_4,25_=9.0, p=0.0001), but not days X treatment (F_12, 75_=1.5, p=0.2). On day 4, sham PBS treated, CHI-MN12 treated and CHI-MN12 PBS infused mice significantly lowered the number of error from day 1 (*p<0.05). Sham saline-treated (n=4), Sham saline-treated ζ-stat infused (n=5), CHI saline-treated (n=4); CHI MN12-treated (n=4), CHI MN12-treated ζ-stat-infused (n=7), CHI MN12-treated saline-infused (n=6) were analyzed.

### ζ-stat blocks Barnes maze acquisition by MN12 injured mice

The reestablishment of L-LTP in injured MN12-treated mice suggests that increased PKMζ expression may also be needed to acquire Barnes maze. This is tested by infusing ζ-stat or PBS bilaterally into the hippocampus 30 minutes prior to the onset of each day of Barnes maze training (Figure 2B). Latency to find the escape hole has significant effects of time (F_2.37, 58.93_=21.3, p<0.0001), treatment (F_5,24_=7.7, p=0.0002) and time*treatment (F_15,72_=2.0, p=0.03). Latency between days 1 and 4 is significantly lowered by sham-injured mice that are saline-treated and infused with PBS (p=0.04), as are injured mice that are MN12-treated (p=0.05) and injured mice that are MN12-treated and infused with PBS (p=0.02) (Figure 2C). In contrast, latency did not significantly differ between days 1 and 4 in injured, saline-treated mice (p=0.5), injured mice that are MN12-treated and infused with ζ-stat (p=0.7), or sham saline-treated mice that are infused with ζ-stat (p=0.9). These data suggest that MN12 lowers Barnes maze latency in injured mice by increasing PKMζ expression.

Errors also show significant effects of time (F_2.37, 58.93_=23.5, p<0.0001) and treatment (F_5,24_=9.0, p<0.0001), but not time*treatment (F_15,72_=1.7, p=0.06). Sham-injured saline-treated mice (p=0.05), injured and MN12-treated mice (p=0.04) or injured mice that are MN12-treated and PBS-infused (p=0.001) show significantly lowered errors between days 1 and 4 (Figure 2D). In contrast the error number does not significantly differ between days 1 and 4 in sham-injured saline-treated and ζ-stat-infused mice (p=0.5), injured saline-treated mice (p=0.8) or injured MN12-treated and ζ-stat-infused mice (p=0.1). These data suggest that MN12 lowered error number by increasing PKMζ expression.

Speed on training day 1 did not differ among the groups ([in meters/min], Sham PBS-treated, 2.52±0.2; sham PBS-treated ζ-stat infused, 3.0 ± 0.3; CHI PBS-treated, 2.4±0.3; CHI MN12 treated 2.6±0.4; CHI MN12-treated PBS infused, 2.9±0.3; CHI MN12-treated ζ-stat infused 2.9 ± 0.3; F_5,114_=0.51, p=0.7). These data suggest similar motor ability in all groups.

## Discussion

This study shows that the injured ipsilesional hippocampus lowers PKMζ in both CA3 and CA1 and lacks both E-LTP and L-LTP (Figure 1, Figure 2A). The contralesional hippocampus lowers PKMζ in CA1 and lacks L-LTP (Figure 1, Figure 2A). Injured mice do not acquire proficiency in Barnes maze (Figure 2C). MN12 treatment bilaterally restores CA1 PKMζ expression, L-LTP and Barnes maze acquisition. Studies using ζ-stat suggest that increased PKMζ underlies restoration of both L-LTP and Barnes maze acquisition. ζ-stat blocks PKMζ action by preventing its binding and subsequent stabilization by KIBRA (19). MN12 may directly increase PKMζ expression or indirectly increase expression by promoting the stabilization of PKMζ by KIBRA. Reduced KIBRA expression has been implicated in cognitive deficits in a murine tauopathy model (26).

MN12 has no effect on CA3 PKMζ expression yet increases expression in CA1 (Figure 1). MN12 likely restores of L-LTP by increasing PKMζ CA1 expression since post-synaptic CA1 PKMζ activity is needed for L-LTP of Schaffer collateral/commissural-CA1 synapses (27-29).

Persistent increases in synaptic plasticity as modeled by L-LTP are needed to acquire hippocampus-dependent spatial tasks such as Barnes maze (2, 3). L-LTP requires increased CA1 PKMζ expression and lowering of PKMζ expression likely contributes to the LTP and Barnes maze deficits. Furthermore, increased PKMζ expression by MN12 has a key role in restorating L-LTP and Barnes maze in injured mice (Figures 1A, 1B, 2C, 2D).

MN12 also restores E-LTP in the injured ipsilesional hippocampus (Figures 1A, 1B). E-LTP is established in the presence of PKMζ inhibitors suggesting that MN12 targets additional processes independent of PKMζ to restore E-LTP in the injured hippocampus.

This study examined the MINO and NAC combination, so it remains uncertain if the individual drugs restore L-LTP or Barnes maze acquisition. Barnes maze acquisition is restored in injured mice with a first dose of MINO, but not NAC at 12 hours-post-injury (1). This suggests that MINO may be sufficient to reestablish PKMζ expression, LTP, and Barnes maze after injury. Dosing MINO beginning 12 hours post-CHI improved Barnes maze acquisition but did not prevent lowered expression of the dendritic marker MAP2 in the ipsilesional hippocampus (1). This study has the caveat that only males were studied yet recent data suggests that MN12 also restores Barnes maze acquisition in injured female mice (Nikulina, Bergold, unpublished results).

### Transparency, Rigor and Reproducibility Statement

This study has been registered at https://osf.io/sqpzf. An analysis plan was not formally pre-registered. Mice were randomly assigned to groups using the Random Picker number generator. Investigators who administered the therapeutic intervention were blinded to group assignment since the MINO plus NAC and saline treatments had an identical appearance. MINO plus NAC was first dosed at 8 hours, 2 and 3 was based upon the previous studies of Sangobowale, et al. (1). Purity of pharmacological reagents was based upon the manufacturer’s certificates of analysis. The specificity of the PKMζ antibody and ζ-stat reagents were verified by the Sacktor laboratory were verified in a recently published study (19). Minocycline is labile in solution, so MINO plus NAC solutions are prepared immediately before use. The PKMζ antibody and ζ-stat peptide used in this study are available upon request from the Sacktor laboratory. This study uses a closed head injury model that is an established standard in the field. The immunofluorescence, long-term potentiation and Barnes maze testing are established outcomes to test efficacy after experimental TBI. Validation of these methods have been confirmed in previous studies from the Sacktor and Bergold laboratories (1, 19). The statistical tests used were based on the assumptions of normal distributions for some parametric tests. Statistical analysis was performed by Dr. Changchi Hsieh, who received a M.A. degree in statistics from Columbia University. The authors provided the full content of the manuscript on BioRxiv as of 4/19/24. For the PKMζ expression study (figure 1) a sample size of 5 mice per group was planned based on an expected effect size of 0.57, calculated to yield 81% power to detect a post-hoc difference between the sham-injured group and the experimental group with largest change in PKMζ expression using 2-way ANOVA with a p-value < 0.05. For the LTP studies (Figures 2 and 4) a sample size of 4 mice per group was planned based upon an effect size of 0.81 calculated to yield 85% power to detect a post-hoc difference between the sham-injured group and the experimental group with largest change in fEPSP at 165 minutes post-HFS during repeated values 2-way ANOVA with a p-value of p<0.05. For the Barnes maze study (figure 4) a sample size of 4 mice per group was based upon an effect size of 0.96 calculated to yield 85% power to detect a post-hoc difference between the sham-injured group and the experimental group with largest change in latency on training day 4 using repeated values 2-way ANOVA with a p-value of p<0.05.

This study used 96 mice. Six injured mice are excluded since time to regain righting reflex was less than 330 seconds. Five mice died within 2 days post-injury due to hemorrhage. Eighteen mice are excluded because one of the two infusion cannulas was placed dorsal to the hippocampus.

